# New Synapse Detection in the Whole-Brain Connectome of *Drosophila*

**DOI:** 10.1101/2025.07.11.664377

**Authors:** Szi-Chieh Yu, J. Alexander Bae, Arie Matsliah, Sven Dorkenwald, Jay Gager, James Hebditch, Ben Silverman, Kyle Patrick Willie, Ryan Willie, Austin T Burke, Thomas Macrina, Sebastian Seung, Mala Murthy

## Abstract

The FlyWire *Drosophila* brain connectome^1^ is a graph of roughly 140K neurons and >50 million synaptic connections^2,3^ reconstructed from the FAFB EM dataset^4^. Challenges in synapse detection were identified for neurons with features such as dark cytosols, axo-axonic synapses, and complex polyadic synapses, due to limitations in ground truth data for these cells and the inherent complexity of these synapse types. To address these issues, we trained new neural networks using iteratively generated ground truth annotations and detected synapses across the entire FAFB dataset, producing what we refer to here as the ‘Princeton synapses.’ These synapses were evaluated in both control regions, such as subareas of the mushroom body calyx and lateral horn, which were also chosen by Buhmann et al.^3^ for evaluation, as well as challenging regions, including Johnston’s Organ neurons (JONs), photoreceptors, and other cell types. The new model shows significant improvements, achieving up to a 0.23 F-score increase in challenging areas, while maintaining performance in control regions. Princeton synapses also show an 8–9% improvement in neuron clustering within cell types and better left/right symmetry scores, especially for photoreceptors. Additionally, neuron type membership can be predicted from connectivity patterns alone with weighted F-scores of 0.93 for Princeton synapses versus 0.91 for Buhmann synapses. The updated Princeton synapses are now accessible via Codex (codex.flywire.ai).

## Synapse Detection Pipeline

The Princeton synapses (detected in the FlyWire/FAFB dataset^1,4^) were generated by a two-step process, modified from the automated method used to generate mouse cortical connectivity^5–7^. Unlike mammalian synapses, *Drosophila* synapses are polyadic, meaning a single pre-synapse can have multiple postsynaptic partners. Therefore, we first detected the postsynaptic terminals, and then the presynaptic and postsynaptic partners were assigned for each detected postsynaptic terminal (Figure 1). For postsynaptic terminal detection, a version of residual symmetric U-Net^8,9^ is trained to detect the postsynaptic terminals, labeled at the postsynaptic sites roughly along the cleft. The model was trained with 150.8 μm3 (7 cutouts) in CREMI-B (https://cremi.org) and 385.88 μm3 (130 cutouts) in FAFB. For validation, 21.47 μm3 (1 cutout) in CREMI-B and 6.04 μm3 (3 cutouts) in FAFB were used. For synaptic partner assignment, a different network was trained to detect the presynaptic and postsynaptic mask. The cell segments with the highest mean probability for presynaptic and postsynaptic masks are assigned as the presynaptic and postsynaptic cells, respectively.

**Figure 1.**
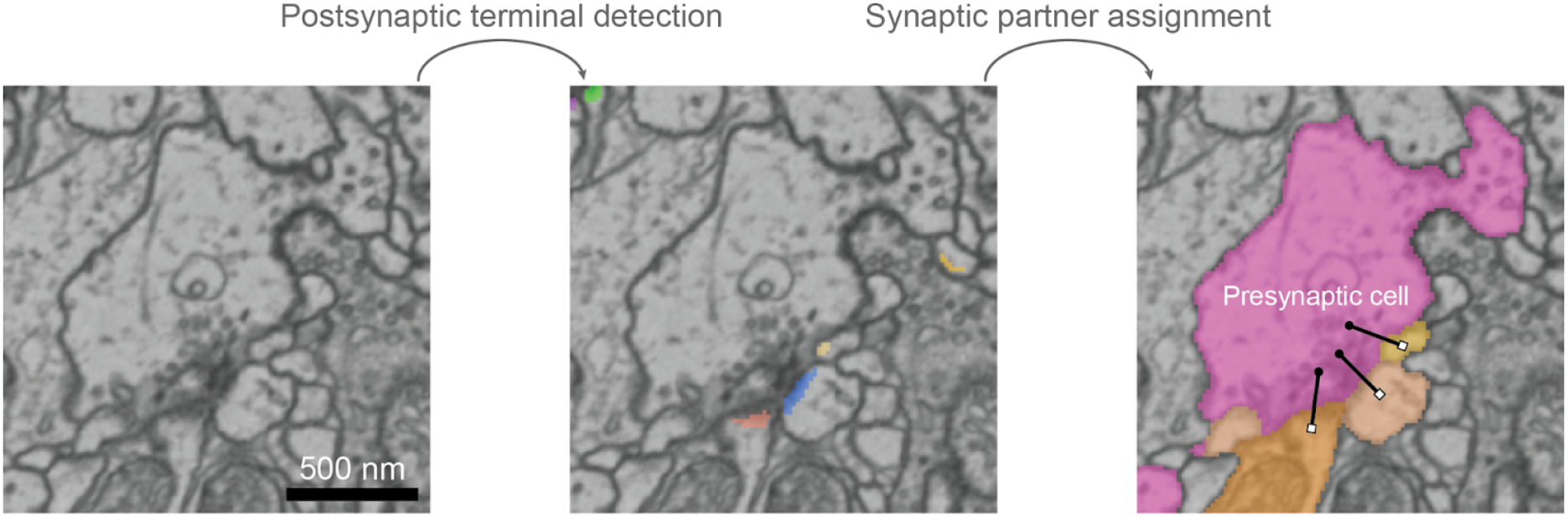
Synapse detection pipeline: Postsynaptic terminal detection processes raw EM images as input and generates a probability map indicating the likelihood of each pixel belonging to the postsynaptic terminal. The model was trained on a resolution of 8 × 8 × 40 nm^3^ with input patch size of 128 × 128 × 20. The predicted output was downsampled by a factor of 2 then thresholded with a pixel value of 0.12. The postsynaptic terminal segments were generated by finding the connected components with 26-connectivity (the voxels are considered connected if they have any contact in any direction, including diagonals, with at least 26 surrounding voxels). The detected terminal may overlap with multiple cells, so the segmented terminal was divided based on the cell segments to ensure each terminal corresponds to a single cell segment. Postsynaptic terminal segments less than 5 voxels in 16 × 16 × 40 nm^3^ were removed.

Synaptic partner assignment predicts two outputs from the raw EM image centered around the detected postsynaptic terminal: the probability of each pixel belonging to either the presynaptic or postsynaptic cell. The assignment model was trained on a resolution of 16 × 16 × 40 nm^3^ with input patch size of 24 × 24 × 8. The assignment model assumes a single presynaptic cell and single postsynaptic cell for each individual postsynaptic terminal. To prevent overlapping assignment, two detected postsynaptic terminals located within 200 nm (based on their centroids) are merged into one terminal and then partners are assigned.

The results can be visualized by line annotations where the end points represent the locations of presynaptic and postsynaptic sites (Figure 1, right). Once the presynaptic partner of the postsynaptic terminal (and cell) is determined within the patch, the closest voxel to the cell segment from any point in the postsynaptic terminal segment is determined. Then, the patch is redrawn, centered around the identified coordinate, and the mean coordinates of voxels that belong to the intended cell segment is computed. Lastly, the algorithm determines the location coordinate by taking the closest voxel of the cell segments from the mean coordinates of both presynaptic and postsynaptic segments.

The final results are stored in DataFrame format. Please see the details below for the meaning of each column.

| Name | Contents |
| --- | --- |
| cleft_segid | Postsynaptic segment id |
| centroid_x, y, z | Centroid coordinates of postsynaptic terminal in $16 \times 16 \times 40 \text{ nm}^3$ resolution |
| bbox_bx, by, bz | Start points of bounding box surrounding the postsynaptic terminal in $16 \times 16 \times 40 \text{ nm}^3$ resolution |
| bbox_ex, ey, ez | End points of bounding box surrounding the postsynaptic terminal in $16 \times 16 \times 40 \text{ nm}^3$ resolution |
| presyn_segid | Presynaptic cell id |
| postsyn_segid | Postsynaptic cell id |
| presyn_x, y, z | Location coordinates of presynaptic cell in 16 x 16 x 40 nm<sup>3</sup> resolution |
| postsyn_x, y, z | Location coordinates of postsynaptic cell in 16 x 16 x 40 nm<sup>3</sup> resolution |
| size | Size of postsynaptic terminal in voxels |

### Iterative Improvement of Synapse Detection Through Identification of Challenging Regions

After the first model was created, we employed an iterative process to enhance our synapse detection model by focusing on regions where prediction from^3^ performed poorly. The initial step involved identifying these challenging regions using several methods:

1. **Community Feedback**: We solicited suggestions from the FlyWire community for areas with suboptimal synapse detection in Buhmann et al.’s model. Each suggestion was carefully reviewed, and regions exhibiting the poorest predictions from the review were selected for further analysis and improvement. These regions included, but were not limited to:
  - Lamina Monopolar Cells
  - HS Cells and DNp20 in the superior posterior slope (SPS) and inferior posterior slope (IPS) regions
  - Cells with dark cytosol in the SPS and IPS
  - DNx01 and DNp15
  - Johnston’s Organ Neurons mechanoreceptor neurons (JONs)
  - Photoreceptors
  - Antennal Motor Neurons
  - DM3_adPN with dark cytosol
  - HRN_VP5 with dark cytosol
  - OA-VPM1~4
  - APL and DPM
2. **Synapse Count Discrepancies Between Hemibrain and FlyWire Brain**: We analyzed neurons with significantly more synapses in the hemibrain dataset than in the FAFB dataset^1^. This analysis was conducted separately for postsynaptic (inputs) and presynaptic (outputs) connections.
  - Postsynaptic Discrepancies: Neurons examined included lLN2F_b, EPG, various vDelta neurons, FB4 neurons (FB4A, FB4D-J), PAM08, PFL3, PFNm, and PFNp.
  - Presynaptic Discrepancies: Neurons included Lobula Columnar Neurons (e.g., LC10, LC17, LC4, LC12), Delta7, LLPC2 neurons (LLPC2a-d), and EPG. VCNs (visual centrifugal neurons) and VPNs (visual projection neurons) from above were reviewed but assigned lower priority due to truncation in the hemibrain dataset.
3. **Underrepresented Synapse Types**: We targeted regions containing neurons that release different peptides or neurotransmitters, recognizing that synapses of these types might be underrepresented in the ground truth data used to train the Buhmann et al. model. We reviewed cells releasing various peptides or neurotransmitters, including Allatostatin-a, Corazonin, NPF, SIFamide, histamine, IPC, DMS, ITP, Hugin-PC, Allatostatin-c, and Capa.
4. **Volume-to-Synapse Count Ratios**: We reviewed neurons exhibiting unusually large ratios of neuronal volume to output synapse count. A disproportionately large volume with relatively few synapses could indicate areas where synapse predictions were insufficient relative to the neuron’s size. A variety of Descending Neurons (DNb06, DNg13, DNpe017, DNge079, DNg26, DNp13, DNg31, DNg39) appeared on our high volume-to-synapse ratio list. This observation is expected because descending neurons do not have axons in the FAFB dataset, contributing to their large volumes with relatively few output synapses. Additionally, we examined single-segment IDs from the following neuron types: Lawf1, Lawf2, Lobula Plate Tangential Cell, Vertical System (LPTC VS), T1, AMMC-A1, AMMC-B1, Medial Ocellar Retinula Cell, Proboscis Motor Neuron, Dm3, BM_InOm, Lat, and a variety of Ascending Neurons (ANs) and other types (CB0804, CB0901, cL19, cLP01, cLP02, cM08a).
5. **Axo-Axonic Synapses**: We targeted regions containing robust axo-axonic synapses to identify areas where the Buhmann et al. model exhibited poor synapse predictions. Axo-axonic synapses are particularly challenging for detection models due to their complex connectivity patterns (please see “Current Limitations” section). The cell types and regions we reviewed included:
  - Mushroom Body Output Neurons (MBONs)
  - CB0687
  - Kenyon Cells
  - LC10 Neurons
  - Lateral Horn
  - Antennal Lobe
  - Dorsal Protocerebrum
  - Antennal Mechanosensory and Motor Center (AMMC)

Upon testing the above cells and regions to identify areas of poor synapse detection in the Buhmann et al. model, we created new ground truth annotations in regions with the worst results (the postsynaptic terminal detection model was trained on cutouts with 2735 synapses in FAFB; the model was validated on cutouts with 111 synapses in FAFB). After training, we qualitatively reviewed the results to identify false positives and false negatives. We then added more ground truth annotations in areas where our model did not perform well and retrained the model again. This iterative process allowed us to progressively enhance the model’s accuracy by targeting specific areas where it previously underperformed.

We reevaluated the model’s performance using additional test volumes related to the issues identified in points 1-5 but located in completely different regions, serving as our final evaluation (see “Evaluation on Synaptic Detection in Specific Cutouts”).

### Ground Truth for Synaptic Partner Assignment

Thirty-nine ground truth cutouts among the cutouts created for postsynaptic terminal detection (see above) were randomly selected for training the pre- and post-synaptic partner assignment model (the cutouts varied in size from 128×128×16 to 512×512×16) in 8 nm resolution). To ensure that the model is able to handle unusual cases, the ground truth volumes included axo-axonic synapses and synapses with the pre- and postsynaptic partners aligned along the z-plane, which was expected to be more challenging compared to the conventional examples.

### Limitations of New Synapse Detection

1. **Clustered axo-axonic synapses**: With our current algorithm, we are not able to capture these synapses since our pipeline assumes a single presynaptic object per detected postsynaptic terminal. For example, only one synapse would be assigned for the magenta postsynaptic terminal (Figure 2a; left) while two presynaptic cells should have been assigned (Figure 2a; right).
2. **Autapses**: Autapses can be found in the *Drosophila* brain^10^. However, while our current method can detect autapses, it cannot reliably distinguish true autapses from erroneous cases. In particular, misassignment can occur that may result in both presynaptic and postsynaptic partner being attributed to the same cell segment, there could be an error in assignment for typical synapse resulting in same presynaptic and postsynaptic cell segment, which may be mistaken for an autapse. As a result, it remains challenging to differentiate real autapses.

**Figure 2.**
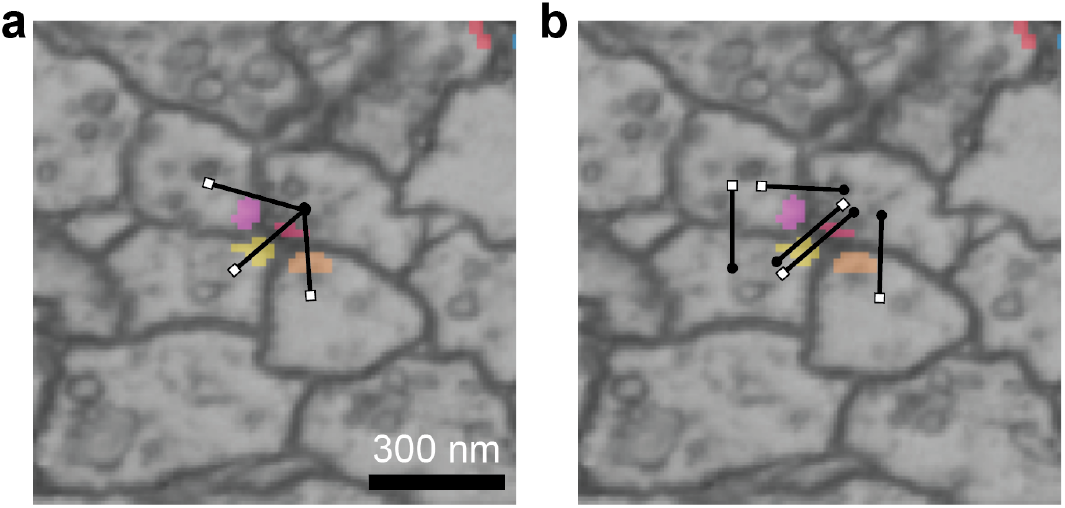
Limitations of new synapse detection: a) predicted synaptic pairs and b) ground truth synaptic pairs for axo-axonic synapses. White square circle: postsynaptic partner, Black circle: presynaptic partner.

### Evaluation of the New Synapse Detection Model

#### Evaluation of 1000 Randomly Predicted Synapses

1000 predicted synapses were randomly selected from diverse regions across the FAFB dataset and evaluated by expert annotators. The evaluated synapses were classified into three categories: correct synapse pairs (90.4%), incorrect postsynaptic terminal detections or assignments (7.3%), and uncertain cases (2.3%) where the validity of the synapse could not be conclusively determined by the experts. This performance was compared with previous predictions by^3^, which had been adjusted for the new alignment of the FAFB EM data used in the FlyWire brain connectome^11^. The comparison highlighted a significant improvement in our current model: whereas the FlyWire Brain connectome reported only 61.6% accuracy^11^, our updated Princeton synapse predictions achieved a much higher precision, validating the model’s enhanced capability to generalize and identify correct synaptic pairs with greater reliability.

### Evaluation of Synaptic Detection on Diverse Test Volumes

To assess the generalization capability and performance of our improved model, we analyzed 16 volumes, each measuring 2048 × 2048 × 1280 nm^3^. These volumes were primarily selected from the challenging cells and areas identified earlier, but were located in entirely distinct areas. In addition, we included completely different cell types and areas where synapse predictions by Buhmann et al. were suboptimal, in order to evaluate the performance of our model on previously “unseen” problematic areas^3^.

12 volumes were selected as **target** volumes where each selected volume contained partial structures from neurons of interest along with parts of their neighboring cells. The primary neurons included were CCHa2R-RA (CB0991) neurons; regions of the mushroom body pedunculus and alpha lobe; vDelta neurons; EPG neurons; Dm9; DmDRA1; Mi1; and various cells characterized by dark cytoplasm, such as R8, JON, DM3_adPN, and L2. Although neighboring neurons are present within these volumes, we refer to each volume by the name of its primary cell type noted here (Figure 3).

**Figure 3:**
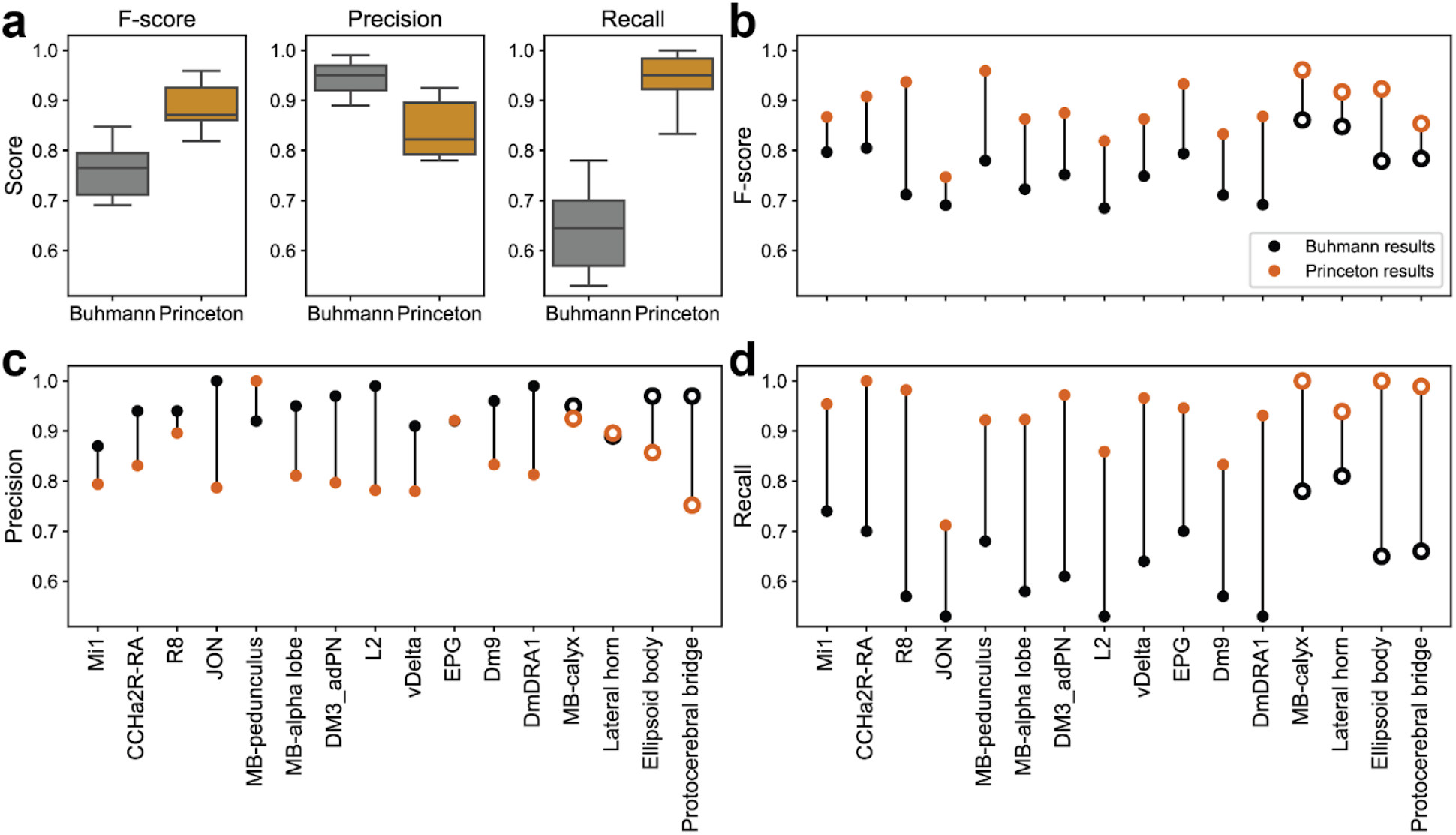
Evaluation results on test cutouts. a) Summary plots showing F-score (left), precision (middle), and recall (right) of filtered Buhmann ^3^ and Princeton (orange) synapse predictions on test cutouts. b-d) F-score (b), precision (c), and recall (d) values for individual test cutout, comparing filtered Buhmann ^3^ and Princeton synapse predictions. The test cutouts consist of 12 target regions (filled) and 4 control regions (open).

Additionally, we examined four **control** volumes from the mushroom body calyx, lateral horn, ellipsoid body, and protocerebral bridge (Figure 3). These regions were randomly selected within the neuropils previously evaluated^3^.

We then generated ground truth synapse annotations in these volumes using line annotations. To ensure an efficient evaluation process, we automatically compared the predicted synapses against the ground truths by first matching synapse segment ID pairs and then searching for line annotations within a 500 nm radius of the ground truth synapses. We adopted this method because the locations of synapse line annotations can vary between different predicted models, even when they refer to the same synapse in the electron microscopy (EM) data. A few synapses near the edges of the dataset might be inconsistently included due to the placement of line annotations, but we consider their number minimal and treat them as noise in our calculations.

The synapse data from^3^ used for comparison were filtered to enhance its accuracy. Synapses with a cleft score ≤ 50 were removed^2^. Additionally, synaptic connections between the same neurons were combined when their presynaptic locations were within 100 nm to reduce falsely redundant synapses between the same neuron pair. Notably, synaptic connections within the same segments had already been filtered according to^3^.

F-scores (containing information on precision and recall) of synaptic predictions across both target and control volumes from filtered Buhmann synapses and Princeton synapses are shown in Figure 3.

In the control regions, including the mushroom body (MB) calyx, lateral horn, ellipsoid body, and protocerebral bridge, the Princeton synapse model exhibited performance comparable to that of the filtered Buhmann synapses. The differences in F-scores between the two models in these regions were similar or favored the Princeton model, indicating it maintains high accuracy in areas where the filtered Buhmann synapse model already performs well. The largest positive F-score difference observed in favor of the Princeton synapses is 0.144, while the smallest is 0.07.

In the target regions, which include challenging cells or areas such as CCHa2R-RA (CB0991), R8, JON, MB-pedunculus, MB-alpha lobe, DM3_adPN, vDelta neurons, EPG, Dm9, and DmDRA1, the Princeton synapse model either shows notable improvements or comparable results over the filtered Buhmann synapses. The largest F-score improvement was observed in the R8 region with a delta F-score of 0.23, while the smallest improvements were in JON, showing a delta F-score of 0.06. These results demonstrate that the Princeton synapse model provides substantial improvement in the more challenging regions and indicates that the model generalizes well across diverse and complex neuronal structures.

When examining precision and recall, Princeton synapses show a significant increase in recall for every volume analyzed (Figure 3). This improvement in recall suggests that the Princeton model is better at identifying true synapses, increasing the likelihood of detecting a larger number of synaptic connections that may have been missed by the Buhmann model. However, this comes with the potential downside of increasing the number of false positives. Despite this, the number of false positives appears to be relatively low. This suggests that while there may be some increase in false positives due to the heightened recall, it does not significantly impact the overall performance of the Princeton synapse model in most regions. A region-specific filter could potentially be applied to further reduce the number of false positives, enhancing the precision of the synapse detection process. However, designing such a filter is beyond the scope of this current analysis and would require further investigation.

### Self Consistency Evaluation with Optic Lobe Cell Types

We used recent results^12^ that reviewed and typed all ~87k cells intrinsic to the optic lobes into 230 cell types (see detailed counts at codex.flywire.ai). Optic-lobe intrinsic neurons provide additional structure, specifically most neurons belong to numerous cell types (100+ cells) and most synapses belong to repeated subcircuits, due to the columnar composition of the compound eye. This structure allows unbiased self-consistency evaluation of the connectivity between cells of given types.

*Connectivity-based similarity of neurons of same type*: For each cell *c*, we defined an output feature vector by the number of output synapses onto neurons of cell type *t*, which runs from 1 to *T*=230. For each cell, we similarly defined an input feature vector by the number of input synapses received from neurons of cell type *t*. The input and output feature vectors were concatenated to form a 2*T*-dimensional feature vector *x*.

As observed previously^12^, cells of the same type are near each other in feature space, while cells of different types are far away. This was quantified using *weighted Jaccard similarity*:

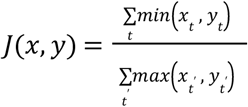

For both Buhmann and Princeton synapses, we calculated the average Jaccard similarity within each cell type to its centroid (mean of all feature vectors in the type cluster). More similar feature vectors within clusters (corresponding to cell types) can be interpreted as more consistent typing.

As depicted in Figure 4, the vast majority of neuron types in both left and right optic lobes cluster slightly better with Princeton synapses. The overall weighted mean similarities to type centroids (across all types) with Buhmann synapses are 0.512 in the left optic lobe and 0.569 in the right optic lobe. With Princeton synapses these are improved to 0.555 in the left optic lobe and to 0.624 in the right optic lobe (8% and 9% improvement respectively).

**Figure 4:**
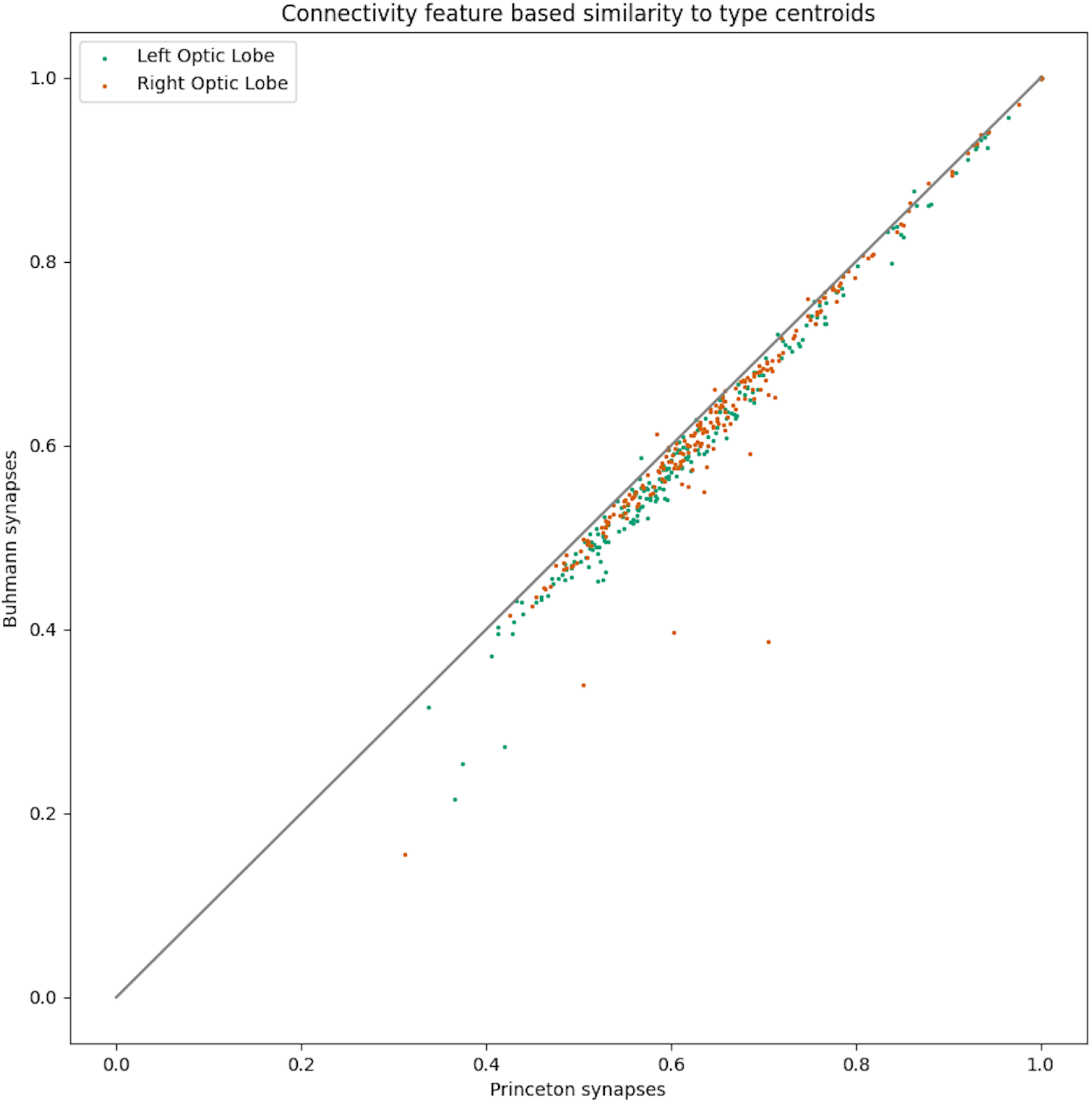
Mean similarity of neurons to their type centroids in the defined feature space with Jaccard similarity metric. Green and orange dots represent types of neurons in the left and right optic lobes respectively. Left and right optic lobes do not intersect (in terms of neurons or synapses), but they contain the same neuron types, and almost the same number of neurons in each type, hence can be thought of as two independent datasets for this comparison. The moderate improvement in clustering within types is consistent in both optic lobes.

When dividing by the average similarity between cluster centroids (Dunn Index) the scores are also slightly improved from 8.92 left / 9.85 right (Buhmann synapses) to 9.23 left / 10.45 right (Princeton synapses).

*Connectivity between types in the left vs right optic lobes*: We also compared the number of synapses between each pair of optic-lobe intrinsic types on the left and right sides of the brain. For any two types, *x* and *y*, we expect the number of synapses from neurons of type *x* to neurons of type *y* to be similar in both hemispheres, due to left/right symmetry. This provides another way to evaluate the consistency of synapse detection: by quantifying how well type-to-type connectivity matches between the two sides of the brain.

The calculation was performed as follows: For cell types *a, b* let *Lsyn(a, b)* denote the total number of synapses where presynaptic/postsynaptic cells are on the left hemisphere and of types a/b respectively. Similarly, let *Rsyn(a, b) denote the same for the right hemisphere. Then:*

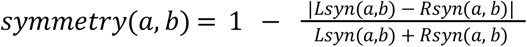

and:

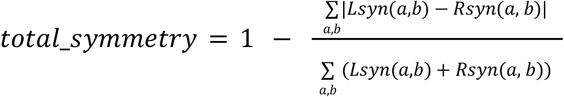

Results

- Buhmann synapses: *total_symmetry = 0*.*91*
- Princeton synapses: *total_symmetry = 0*.*92*
- Percent pairs where Princeton symmetry is better/same/worse: 31% / 52% / 17%

It is worth noting that a more significant improvement in L/R symmetry is evident in the Lamina, specifically for the photoreceptor types (R1-6, R7 and R8). Measuring the same quantities but only for pairs of types where at least one of them is a photoreceptor we get:

- Buhmann synapses: *total_symmetry = 0*.*83*
- Princeton synapses: *total_symmetry = 0*.*88*
- Percent pairs where Princeton symmetry is better/same/worse: 67% / 0% / 33%

*Compact connectivity predicates*: Prior work^12^ found the existence of a set of compact logical predicates based on connectivity that predict type membership of every cell in a type in the optic lobe with high accuracy. For a given neuron, we define the attribute “is connected to input type t” as meaning that the neuron receives at least one connection from some neuron of type t. Similarly, the attribute “is connected to output type t” means that the neuron makes at least one connection onto some neuron of type t. An optimal predicate is constructed for each cell type that consists of 2 sets: input types and output types. Both sets are limited to size 5 at most, and they are optimal with respect to the F-score of their prediction of the subject type, defined as follows:

- **Recall** of a predicate for type T is the ratio of true positive predictions (cells matching the predicate) to the total number of true positives (cells of type T). It measures the predicate’s ability to identify all positive instances of a given type.
- **Precision** is the ratio of true positive predictions (predictions that are indeed of type T) to the total number of positive predictions made by the logical predicate.
- **F-score** is the harmonic mean of precision and recall - a single metric that combines both precision and recall into one value.

The process for computing the predicates is exhaustive - for each type we look for all possible combinations of input type tuples and output type tuples and compute their precision, recall and f-score. For example, the logical predicate “is connected to input type Tm9 and output type Am1 and output type LPi15” predicts T5b cells with 99% precision and 99% recall. For all but three of the previously identified types^12^, a logical predicate with 5 or fewer input / output attributes was found with average f-score 0.91, weighted by the number of cells in type.

Existence of these compact predicates is another consequence of the columnar organization of the cell types in the optic lobes, and the average f-score is a way to quantify self consistency of the connectivity between types. In comparing Buhmann synapses with Princeton synapses, we find that Buhmann synapses weighted average f-score is **0.91**, and Princeton synapses weighted average f-score is **0.93**.

### Breakdown of synapse count differences between Buhmann and Princeton synapses

Overall the Princeton method has detected more synapses compared to Buhmann (76,944,499 vs 54,492,922) and these also make up more connected neuron pairs (22,285,323 vs 15,091,983). Figure 5 shows the breakdown of synapse count differences by region (neuropil), Figure 6 shows the breakdown by presynaptic cell type, and Figure 7 shows the breakdown by post synaptic cell type.

**Figure 5:**
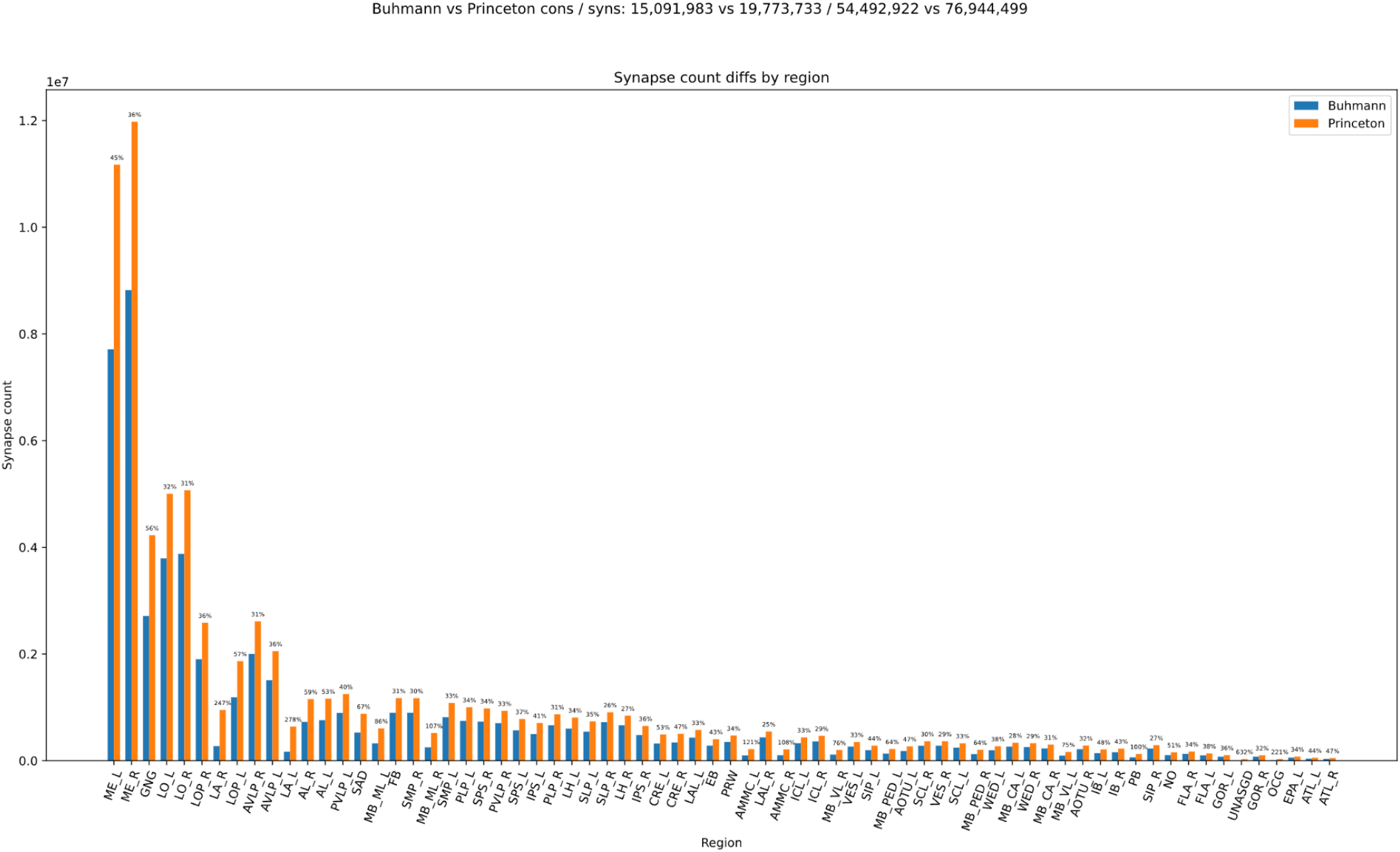
Breakdown of synapse count differences between Buhmann and Princeton synapses by region.

**Figure 6:**
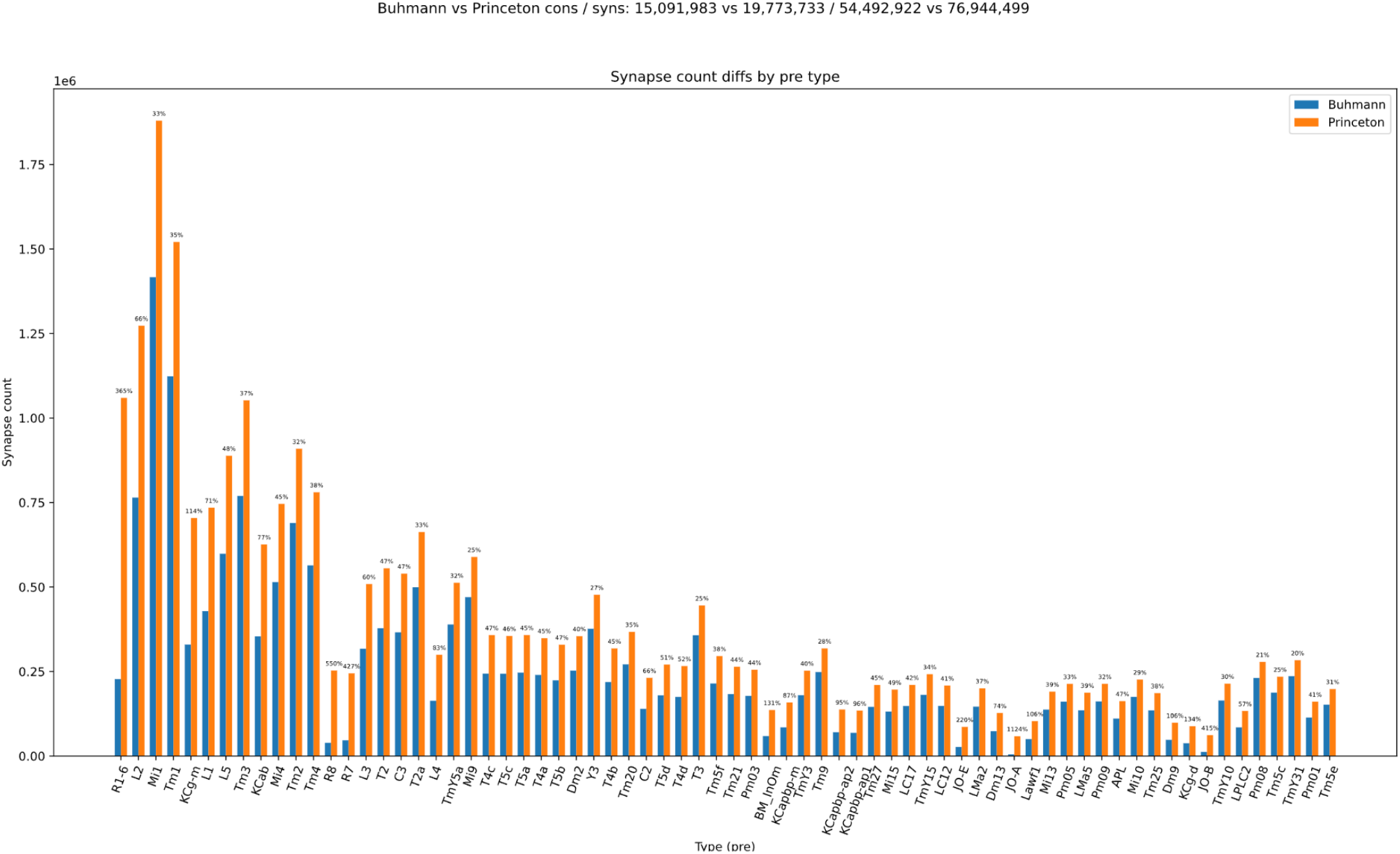
Breakdown of synapse count differences between Buhmann and Princeton synapses by **presynaptic** cell type (top impacted types).

**Figure 7:**
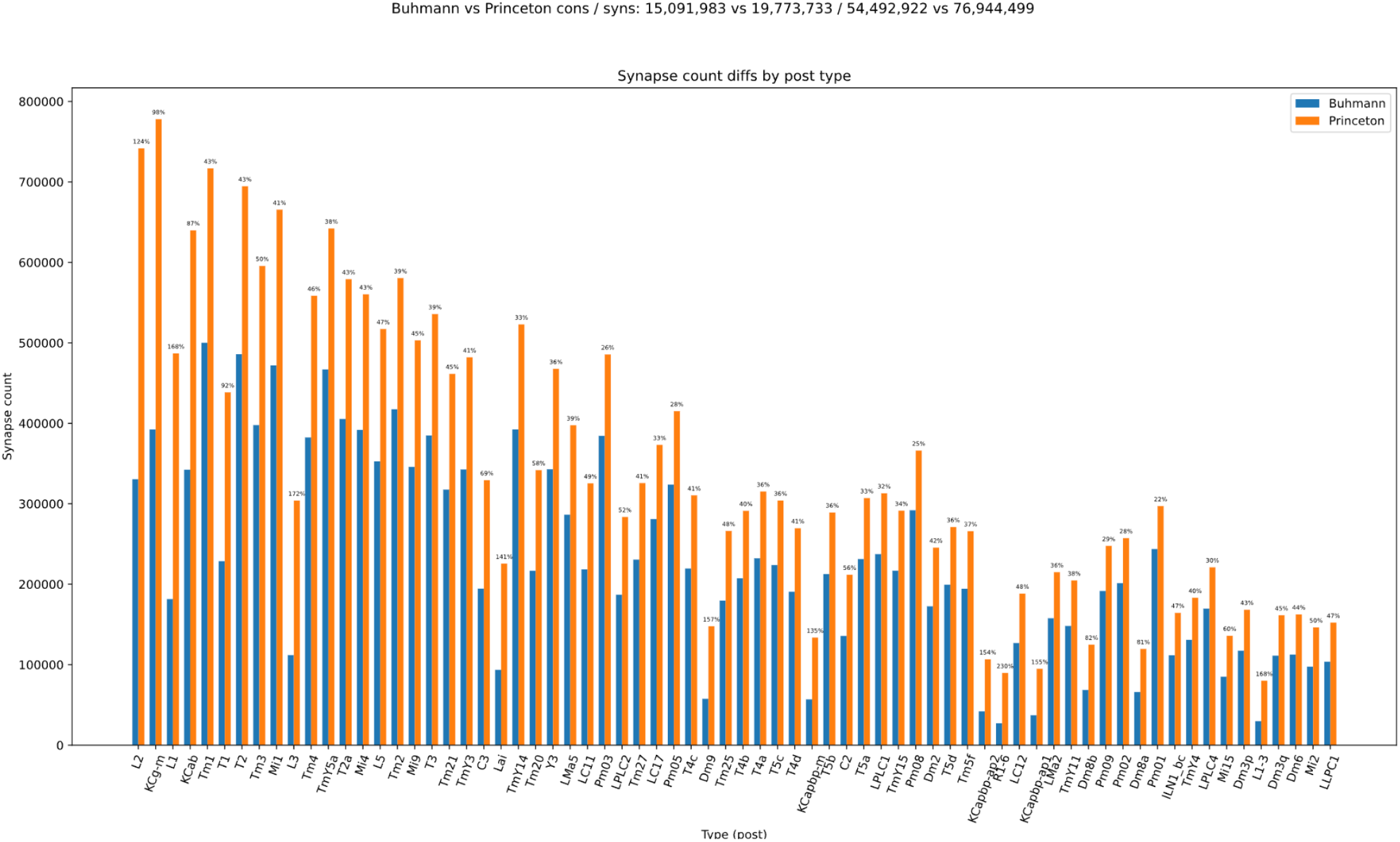
Breakdown of synapse count differences between Buhmann and Princeton synapses by **postsynaptic** cell type (top impacted types).

## Availability of Princeton synapses

The Princeton synapses can be downloaded for analysis from FlyWire Codex (https://codex.flywire.ai/api/download?dataset=fafb). The “Connections Princeton No Threshold” CSV file contains one row for every connected pair of proofread cells and neuropil (from FlyWire FAFB Full Brain snapshot v783). First and second columns contain the FlyWire FAFB Root IDs of the connected pair (‘pre/from’ and ‘post/to’ respectively), the third column contains the neuropil abbreviation, the fourth contains the number of synapses (aggregated across all connection sites of the respective pair and neuropil). FlyWire Brain connectome with Princeton synapses is also available for exploration in Codex (default since July 2025).

Similar to the original Buhmann synapses, the connectome loaded into Codex is thresholded at 5+ synapses per connected pair of neurons.

